# Topoisomerase IB interacts with genome segregation proteins and is involved in multipartite genome maintenance in *Deinococcus radiodurans*

**DOI:** 10.1101/2020.06.24.169326

**Authors:** Swathi Kota, Reema Chaudhary, Shruti Mishra, Hari S. Misra

## Abstract

*Deinococcus radiodurans*, an extremophile, resistant to many abiotic stresses including ionizing radiation, has 2 type I topoisomerases (drTopo IA and drTopo IB) and one type II topoisomerase (DNA gyrase). The role of drTopo IB in guanine quadruplex DNA (G4 DNA) metabolism was shown *in vitro*. Here we report that *D. radiodurans* cells lacking drTopo IB (Δ*topoIB*) show sensitivity to G4 DNA binding drug (NMM) under normal growth conditions. The activity of G4 motif containing promoters like *mutL* and *recQ* was reduced in the presence of NMM in mutant cells. In mutant cells, the percentage of anucleated cells was more while the ploidy numbers of genome elements were less as compared to wild type. Protein-protein interaction studies showed that drTopo IB interacts with genome segregation and DNA replication initiation (DnaA) proteins. The typical patterns of cellular localization of GFP-PprA were affected in the mutant cells. Microscopic examination of *D. radiodurans* cells expressing drTopo IB-RFP showed its localization on nucleoid forming a streak parallel to the old division septum and perpendicular to newly formed septum. These results together suggest the role of drTopo IB in genome maintenance in this bacterium.

## 1. Introduction

DNA transaction processes like unwinding, rewinding, movement of proteins on DNA, the formation of higher-order structures during DNA compaction leads to topological entanglements. These topological barriers if unresolved, pose threats to genome stability. DNA topoisomerases are ubiquitous, found in all organisms which resolve the topological constraints in the DNA. Based on the structure and kind of action topoisomerases are classified into type I and type II enzymes. Unlike type II DNA topoisomerases that cleave both the DNA strands and change the linking number in the multiples of two, the type I enzyme reversibly cleaves one strand at a time and the change the linking number in the increment of one. Type I topoisomerases are subdivided into type IA and type IB subfamilies. Type IA topoisomerases cleave the DNA strand and make a 5’-phosphotyrosine covenant intermediate with a free 3’-OH strand, while type IB topoisomerases prefer double-stranded DNA, forms a 3’-phosphotyrosine covenant intermediate with a free 5’-OH strand. Type IA enzymes relax only negatively supercoiled DNA, while type IB enzymes relax both negative and positive supercoiled DNA. Both the type I enzymes conserves the energy of the phosphodiester bond in the protein-DNA covalent intermediate, thus overall cleavage and religation reactions with DNA strand do not require ATP. Type II topoisomerases are also sub divided into type IIA and type IIB, where type IIA enzymes are present in all domains of life while type IIB enzymes are present in archaea and in some higher plants (Champoux, 2001; Chen et al., 2013). *Escherichia coli* contains 4 topoisomerases, 2 type I topoisomerases (Topo I encoded by *topA*, and Topo III encoded by *topB*) and 2 type II toposiomerases (DNA gyrase and Topo IV) to maintain the steady-state level of supercoiling. Both the type I topoisomerases play an important role in genome maintenance. For instance, Topo I regulates super helicity and R-loop formation and Topo III is involved in chromosome segregation. *E. coli* double mutant of *topB* and *topA* with a compensatory mutation in either *gyrA* or *gyrB* shows unsegregated nucleoids with an unusual morphology, which eventually leads to growth arrest (Zechiedrich et al., 2000; Usongo et al., 2013). In *B. subtilis* DNA gyrase co-localizes with replication machinery and tackles the positive supercoils associated with the DNA replication. TopA is seen in proximity to SMC complex which generally assists in DNA compaction and packaging, and TopoIV is distributed on nucleoid at many places (Tadesse and Graumann, 2006).

In yeast and human type IB topoisomerase (Top I) relaxes positive and negative supercoils generated during the processes of DNA replication, repair, recombination and transcription. In budding yeast, either loss of Top I or increased number of Top I cleavage complexes leads to genome instability. At highly transcribed regions of the genome, an active role of Top I in preventing excessive gross chromosomal rearrangements and, loss of heterogeneity was noticed (Andersen et al., 2015). Topo I was showed to prevent the non-canonical B DNA structures formation like guanine quadruplex DNA (G4 DNA) and RNA-DNA hybrid structures (R-loops) in the non - template strand during transcription (Yadav et al., 2016). In human, the topoisomerases are the main enzymes that are targeted in cancer treatment because of their important role in DNA metabolism (Delgado et al., 2018).

*Deinococcus radiodurans* is extremotolerant bacterium, shows resistance to many abiotic stresses including ionizing radiation. Strong anti-oxidative mechanisms, efficient DNA double-strand break repair pathways along with other features like polyploidy and multipartite genome system are the key features attributed to the ionizing radiation resistant phenotype in this bacterium (Cox and Battista 2005; Misra et al., 2013; Slade and Radman 2011). The Genome sequence of *D. radiodurans* encodes 2 type I topoisomerases such as drTopoIA (DR_1374) and drTopoIB (DR_0690) and one type II topoisomerase (White et al., 1999). The DNA gyrase comprised of GyrA (DR_1913) and GyrB (DR_0906) subunits has been characterized as a single type IIA family topoisomerase in this bacterium (Kota et al., 2016). PprA, a pleiotropic protein involved in radioresistance differentially regulates deinococcal gyrase activities, while *pprA* mutant strain is more sensitive to gyrase inhibitors as compared to wild type (Kota et al., 2016; Devigne et al., 2016). *In vitro* studies have shown functional interaction of drTopo IB with PprA protein, as well as its role in the resolution of G4 DNA structures (Kota et al., 2014a; Kota et al., 2015a). Here, we report the functional interaction of drTopo IB with the elements contributing to genome integrity *in vivo*. We showed that *topoIB* mutant strain is sensitive to G4 DNA binding ligand N-Methyl mesoporphyrin (NMM) and the promoter activity of *groESL, mutL* and *recQ* is down regulated under normal growth conditions under mutant background. Interestingly, drTopo IB interacted with genome partitioning proteins but not with cell division proteins. When the effect drTopo IB deletion on genome maintenance was studied, Δ*topoIB* cells showed a reduced copy number of genome elements and more number of anucleate cells as compared to wild type. The *topoIB* mutant showed the arrest of the typical cellular dynamics of PprA during post-irradiation recovery. The fluorescent-tagged drTopo IB localization on the nucleoid was found to be perpendicular to the plane of cell division, which changed dynamically during different phases of growth. All these results suggested that drTopo IB plays an important role in genome maintenance perhaps by interacting with genome segregation proteins and by regulating the dynamics of G4 DNA structure in *D. radiodurans*.

## 2. Material and methods

### 2.1 Bacterial strains and materials

*Deinococcus radiodurans* ATCC13939 strain was a gift from Professor J Ortner, Germany (Schafer et al., 2000). Wild type D. radiodurans and their derivatives were grown aerobically in the medium TGY (0.5% Bacto tryptone, 0.3% Bacto yeast extract, 0.1% glucose) broth at shaking speed of 180 rpm or on agar plates if required, at 32°C with appropriate antibiotics. *E. coli* strain NovaBlue was used for recombinant plasmids maintenance, while *E. coli* strain BTH 101 was used for protein-protein interaction studies. All the recombinant techniques used in this study were as described in (Green and Sambrook, 2012). For *E. coli*, antibiotics kanamycin (25 or 50 µg/ml), ampicillin (100 µg/ml), spectinomycin (100µg/ml) whereas for *D. radiodurans* kanamycin (10 µg/ml), chloramphenicol (5 µg/ml), spectinomycin (70 µg/ml) were used as mentioned in the respective parentheses to maintain the recombinant constructs.

### 2.2 Generation of disruption mutant of drTopo IB of *Deinococcus radiodurans*

Genomic sequence ∼800 bp downstream and upstream to the drTopo IB encoding sequence was PCR amplified with sequence specific primers (Table S1) from the genomic DNA and cloned in the vector pNOKOUT (Khairnar *et al*., 2008) to yield pNOKtopoIB. The recombinant plasmid was linearized with *Sca*I and transformed into *D. radiodurans*. The transformants were plated on TGY plates supplemented with kanamycin (10 µg/ml). Visible colonies obtained were grown for a number of generations in TGY broth supplemented with kanamycin (10 µg/ml) to obtain the homozygous *topoIB* disruption mutant. Homozygous replacement of *topoIB* with *nptII* in the whole genome was confirmed by PCR amplification using gene specific primers (Table S1). Similarly, *nptII* positive clones were screened for absence of *topoIB* coding sequence using the internal primers (Table S1). The clones showing the absence of *topoIB* internal region were considered homozygous disruption mutants of *drtopoIB* (hereafter referred as *topoIB cells*) (Fig. S1).

### 2.3 Construction of expression plasmids

The drTopo IB coding sequence (1041bp) was PCR amplified from *D. radiodurans* genomic DNA using specific primers (Table S1) and was cloned under *groESL* promoter in pRadgro (Misra *et al*., 2006). The recombinant plasmid was transformed in *D. radiodurans* wild type and *topoIB* cells. The drTopoIB-RFP fusion was constructed by cloning the coding sequence along with the histidine tag from pETtopoIB plasmid (*his-topoIB*) and cloned upstream of red fluorescent protein coding region in pDsRed plasmid to yield pDsRedtopoIB (Kota et al., 2014a). The *his-topoIB-rfp* region was PCR amplified from pDsRedtopoIB plasmid and cloned in pRadgro plasmid and transformed in *D. radiodurans* cells. Similarly, *his-topoIB* was also cloned in pRadgro plasmid yielding pRadHistopo. Transformed cells were checked for the expression of His-tagged protein using antibodies against polyhistidine tag (Fig S2). For protein-protein interaction studies using bacterial two hybrid system and co-immunoprecipitation, the drTopo IB coding sequence was cloned in bacterial two hybrid vectors pKNT25, pUT18 and the expression of T18 and T25 tagged drTopo IB fusion in *E. coli* was checked by western blotting (Fig S2). Various proteins involved in genome segregation, cell division and DNA replication initiation protein (DnaA) encoding sequences were cloned in BTH vectors earlier (Maurya et. al., 2016; Maurya et al., 2019). Similarly, T18 tagged ParB coding sequences were cloned in pVHS559 plasmid earlier (Maurya et al., 2019a) and GFP-PprA, FtsZ-GFP, DivIVA and ParA3 were also constructed earlier (Kota et al., 2014b, Modi et al., 2014; Chaudhary et al.,2020).

### 2.4 NMM treatment and promoter expression analysis

To study the *topoIB* disruption effect on promoter expression, the plasmids containing different deinococcal promoters like pRADZ3 (*groESL* promoter without G4 DNA motif), pRADmutL (*mutL* promoter with G4 DNA motif) and pRADrecQ (*recQ* promoter with G4 DNA motif) were transformed into wild type and *topoIB* cells as described previously (Kota et al., 2015a). The transformed colonies obtained for both wild type and Δ*topoIB* were grown in TYG with and without G4 DNA stabilizing agent 50 nM NMM (Frontier Scientific) at 32°C. The early stationary phase cells of these transformants were treated with gamma radiation wherever necessary at a dose rate 1.81 kGy/h (60Co; Gamma Cell 5000; Board of Radiation and Isotopes Technology, DAE, Mumbai, India). A separate set of cultures kept on ice were considered as unirradiated controls. The β-galactosidase enzymatic activity under different promoters was monitored in wild type and Δ*topoIB* cells as described previously (Meima, et al., 2001).

### 2.5 Protein-protein interaction studies

#### In recombinant *E. coli* host

Interaction of drTopo IB with cognate genome segregation proteins was studied in the heterologous host *E. coli* using Bacterial Two Hybrid System (BACTH) and co-immunoprecipitation (Co-IP) techniques. Briefly for BACTH analysis, the drTopo IB interaction with DnaA (Dr_0002) was studied by co-transforming pUTtopoIB (topoIB-C18) and pKNTDA (DnaA-C25) plasmids in *E. coli* BTH101. Similarly, drTopoIB-C25 and ParB protein encoding sequences in pUT18 plasmid (ParB1-C18, ParB2-C18 and ParB3-C18) were co-transformed in BTH101. Cells co-transformed with empty vectors pUT18 and pKNT25 were used as negative controls and those transformed with plasmids pKNTEFZ (*E. coli* FtsZ) and pUTEFA (*E. coli* FtsA) were used as positive controls. The co-transformants obtained were grown in liquid LB broth containing kanamycin (50 µg/ml), ampicillin (100 µg/ml) and IPTG (0.5 mM) for 24-36h at 30°C and β-galactosidase enzyme activity was assayed and calculated as described earlier in (Maurya et. al., 2016; Battesti and Bouveret, 2012) and plotted using GraphPad Prizm5.

Co-immunoprecipitation was performed in *E. coli* strain BL21 (DE3) pLysS cells expressing His-topoIB and T18 tagged genome segregation proteins in different combinations using the kit Protein G immunoprecipitation from Sigma-Aldrich as described previously (Maurya et. al., 2016). Briefly, the co-transformed cells were sub-cultured in liquid medium with appropriate antibiotics and 0.5 mM IPTG was added when the culture reached to early log phase. The cells were pelleted 3h after the induction and proceeded for the preparation of cell free extract. The cell free extract of the transformed cells was made by suspending the cells in RIPA buffer containing 50 mM Tris-base, 100 mM NaCl, 5 mM EDTA, 0.5% triton-X 100, 1 mM PMSF, 1 mM DTT, 0.3 mg/ml lysozyme and 50 μg/ml protease inhibitor cocktail and incubated for 2h at 4°C on a rotator followed by sonication to lyse the cells. Total proteins obtained after centrifugation were then incubated with antibodies against polyhistidine tag and proteins were immunoprecipitated as per the manufacturer’s protocol. The immunoprecipitate was separated on SDS-PAGE, blotted to PVDF membrane and then hybridized with T18 tag monoclonal antibodies. The alkaline phosphatase conjugated anti-mouse secondary antibodies were used for hybridization and for detecting signals BCIP/NBT substrates (Roche Biochemical, Mannheim) was used.

#### In native host *Deinococcus radiodurans*

Interaction of drTopo IB with different ParB proteins in *D. radiodurans* was checked by co-immunoprecipitation as described in (Maurya et. al. 2018). The *D. radiodurans* cells were transformed with recombinants plasmids harbouring His-TopoIB (pRadHistopo) and T18-ParBs (on pVHS559 plasmid). The colonies obtained were scored on spectinomycin (75 μg/ml) and chloramphenicol (8 µg/ml). Individual colonies were grown in liquid broth overnight in presence of 20 mM IPTG and co-immunoprecipitation was done as described previously (Maurya et. al. 2018).

#### *In silico* studies

For *in silico* protein-protein interaction analysis, the amino acid sequences of deinococcal topoisomerase IB and DnaA proteins were retrieved from uniprotKB (http://www.uniprot.org/) and submitted for 3D model building (http://protein.ict.ac.cn). Stereo chemical refinement of the models for Ramachandran plots was done with ModLoop server. The refined models were validated using Swiss model workspace encompassing the package of Anolea, DFire, QMEAN, Gromos, DSSP, Promotif and ProCheck (http://swissmodel.expasy.org/workspace/). The HADDOCK was chosen for molecular docking and the best-scored dock complex obtained from HADDOCK was selected for visualization by PYMOL or UCSF chimera.

### 2.6 Determination of ploidy using quantitative real-time PCR

The number of copies of different genetic elements in wild type and *topo IB* mutants was determined as described in (Breuert et al., 2006). Briefly equal numbers of cells in appropriate growth stage were harvested and exact cell count was determined with Neubauer cell counter. The cells lysis was performed after washing with 70 % ethanol solution in a buffer containing 20 mM Tris pH 7.6, 1 mM EDTA and 5mg/ml lysozyme at 37°C. The cell lytic efficiency was determined using Neubauer counting chamber and the genomic DNA integrity was analysed by agarose gel electrophoresis. The genomic DNA was serially diluted and used for determining genomic copy number by quantitative Real-Time PCR as described in (Mishra et al., 2019). For standardization of qPCR two individual genes for each replicon with almost same PCR efficiency were taken for copy number determination as given in (Mishra et al., 2019). Briefly, ∼300 bps of DNA fragment of each gene from *D. radiodurans* R1 (ATCC13939) genomic DNA was PCR amplified and purified by Gel Extraction kit (Qiagen, Inc) and amount was calculated using nanodrop. Final concentration was estimated with the molecular mass computed with ‘oligo calc’ (www.basic.northwestern.edu/biotools). Different serial dilutions were generated and used for qPCR standardization. By following MIQE guidelines, the PCR was performed and the optimum cycle threshold (Ct) values were estimated. For each replicon the copy number was estimated by relating the standard curves with test sample Ct values. The genomic copy number per cell for each replicon was calculated from both the genes by estimating the number of cells present at cell lysis. The aaverage number obtained based on calculations from both the genes per replicon was represented with appropriate bio-statistical analysis and plotted.

### 2.7 Microscopic studies

Microscopic studies of *D. radiodurans* and *topoIB* mutant cells and their derivatives were carried out with an inverted fluorescence microscope (Olympus IX83 equipped with an Olympus DP80 CCD monochrome camera) as described in (Maurya et al., 2019b). In general, the exponentially growing cells were harvested and washed with phosphate buffer saline (PBS). The cells were suspended in PBS and stained with DAPI (4’, 6-diamidino-2-phenylindole dihydrochloride) (0.2 μg / ml) for nucleoid and Nile red (1μg / ml) for the membrane as required. The membrane in cells expressing drTopoIB-RFP was stained with vancomycin-fluorescent-BIODPY (Van-FL) as described earlier (Tiyanont et al., 2006). After staining, the cells were washed with PBS for 3 times to remove excess stain and mounted on glass slides coated with 0.8% agarose. The wild type and mutant cells expressing GFP-PprA on pVHpprA were processed as described earlier (Kota et al., 2014b). The stained cells were observed under DAPI, TRITC (tetramethylrhodamine isothiocyanate) and FITC channels for DAPI, Nile red and RFP, and GFP and Van-FL fluorescence, respectively. The inbuilt cell Sens software was used for aligning the images. The images captured under different channels were represented as isolated as well as merged. The Adobe Photoshop 7.0 was used to adjust brightness and contrast of images. Nearly 300 cells from wild-type, *topoIB* mutant and their derivatives were analyzed and data plotted in GraphPad Prizm 5.

## 3. Results

### 3.1 drTopo IB regulates G4 DNA response in *Deinococcus radiodurans*

*Deinococcus radiodurans* genome has numerous G4 DNA forming motifs which were earlier shown to play an important role in gamma radiation responsive gene expression in this bacterium (Mishra et al., 2019). Deinococcal topoisomerase IB (drTopo IB) was also shown previously to resolve *in vitro* the intra molecular G4 DNA structures (Kota et al., 2015b). To study *in vivo* role of drTopo IB in *D. radiodurans* G4 DNA metabolism, Δ*topoIB* mutant was generated and effect of drTopo IB deletion on G4 DNA mediated phenotypes was monitored. Under unirradiated conditions, the *topo IB* cells showed higher sensitivity to NMM with an extended lag phase as compared to wild type. *In trans* expression of topo IB on moderate copy plasmid in *topoIB* cells helped to near complete recovery of wild type growth pattern (Fig. 1). This suggests an *in vivo role* of drTopo IB in G4 DNA metabolism in *Deinococcus radiodurans*.

**Fig 1.**
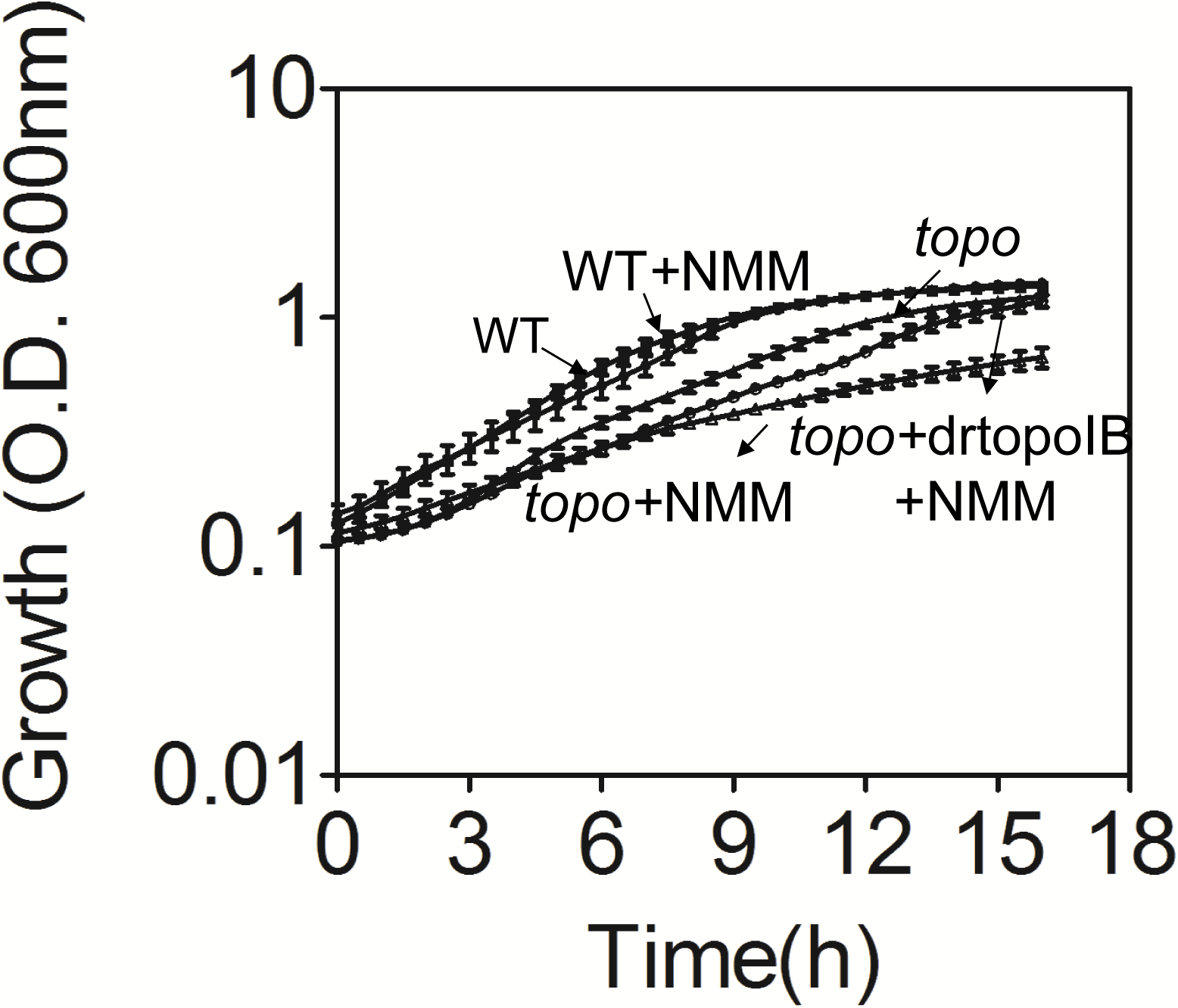
TopoIB role in G4 DNA regulated growth under normal conditions. The *D. radiodurans* R1 wildtype (●) and *topoIB* mutant (▲) cells were grown in TGY and wildtype (■), *topoIB* mutant (△) and *topoIB* cells expressing native His-topoIB (drtopoIB) on plasmid (◒) cells were grown in TGY in presence of 50 nM NMM at 32°C and growth was monitored at 600nm online in a microplate reader (Synergy H1 hybrid multimode).

The promoter activity of three promoters viz; *groESL, mutL* and *recQ* were compared in *topoIB* typical growth conditions and in NMM presence as described in (Kota et al. 2015a). Results showed the downregulation of all three promoters activity in *topoIB* cells in the presence of NMM when compared with cells grown without NMM, which had affected a very little in wild type (Fig 2). The impact of drTopo IB deletion on these promoters activity was still higher when mutant was grown under the influence of both NMM and gamma radiation (Fig 2B, 2C & 2D). Moreover, *groESL* promoter activity also reduced in *topoIB* cells in treatment to NMM at usual growth conditions, whereas in wild type cells the promoter expression levels were similar in NMM and gamma radiation treated/untreated cells (Fig 2A, 2B). Earlier, we have shown that G4 DNA differentially regulates the *mutL* and *recQ* promoter activities after ionizing radiation treatment in wild type cells, depending on the topology of G4 DNA motif present in the promoter (Kota et al. 2015a). This differential effect on G4 DNA containing promoters was not observed in *topo IB* cells (Fig 2C & 2D).

**Fig 2.**
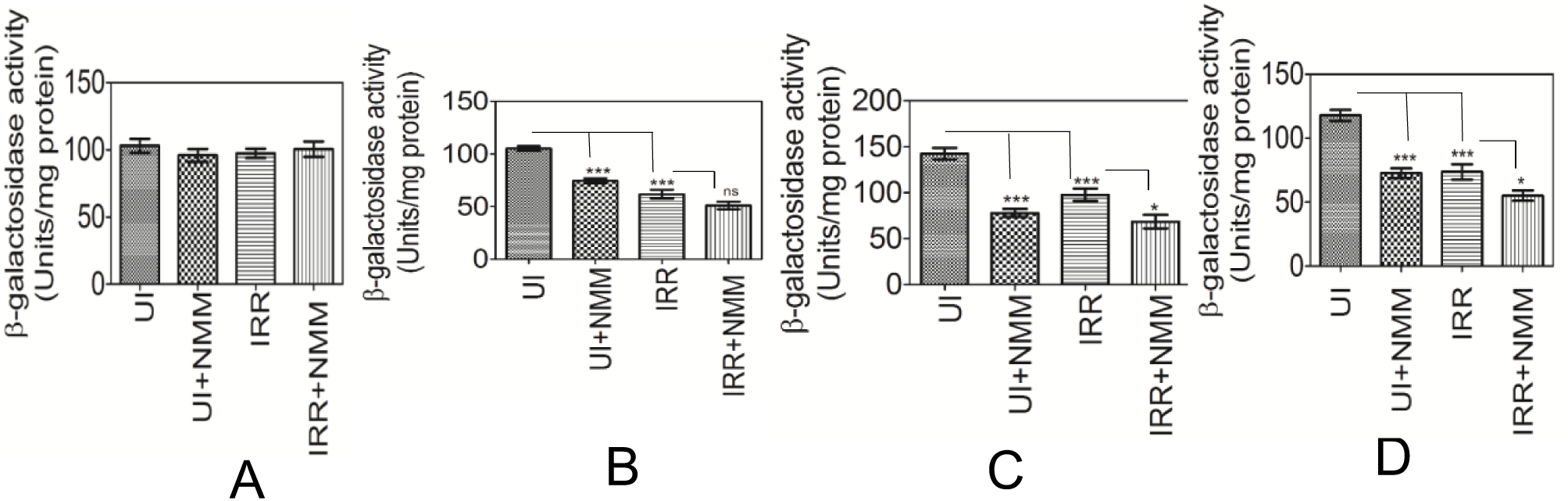
Effect of *topoIB* disruption on promoter activity in *Deinococcus radiodurans*. The wild type *D. radiodurans* (A) and *topoIB* (B, C, D) cells expressing β-galactosidase under the control of *groESL* on pRADZ3 (A, B), *recQ* on pRADrecQ (C), and *mutL* on pRADmutL (D) promoters were grown with (+NMM) and without NMM under normal (UI) and gamma irradiated (IRR) conditions and the levels of β-galactosidase activity was measured as given in materials and methods. The β-galactosidase enzyme activity was shown ± the SD (n=9). The Student’s t test was performed to check the significance of the difference. The P values obtained at 95% confidence intervals were represented as (*) for <0.05, (**) for <0.01 and (***) for <0.001 and ns (not significant).

In bacteria, the role of DNA supercoiling on gene expression has been demonstrated. The supercoiling state of DNA affects gene expression indirectly, by affecting the activity of regulatory proteins (Houdaigui et al., 2019). Topoisomerase IB is known to relax both negative and positive DNA supercoiling (Sissi and Palumbo 2009). The negative supercoiling is shown to favour the formation of non-canonical structures like G4 DNA (Panayotatos and Wells, 1981; Kohwi and Kohwi-Shigematsu 1988; Murchie and Lilley, 1992; <u>Sekibo</u> and <u>Fox</u>, 2017) and if that contributes to the sensitivity of *topoIB* cells to NMM under normal growth conditions cannot be ruled out. Alternatively, drTopo IB was shown to resolve the G4 DNA structures *in vitro* and the absence of this enzyme if has increased the number of G4 DNA structures and that got stabilized by NMM could also explain the higher sensitivity of *topoIB* cells to NMM under normal conditions. But the differential regulatory effect of G4 DNA on promoter activity as observed earlier in wild type cells was not observed in *topo IB* cells and all the promoters responded alike to NMM in the mutant. This suggests that drTopo IB regulates the G4 DNA promoter activity irrespective of the G4 DNA topology. As *groESL* promoter has no G4 DNA motif, the role of drTopo IB in the regulation of promoter activity also seems to be through changing DNA supercoiling.

### 3.2 drTopo IB disruption affects genomic stability in dividing populations

The *topoIB* cells were examined under the microscope and compared with the wild type cells. The deletion of drTopo IB resulted to a significant surge in the frequency of anucleate cells (Fig 3A, 3B) and a substantial number of cells showed the diffused nucleoid as compared to wild type cells (Fig S1). When the copy number of individual genome elements was estimated by qPCR, a significant decrease in copy number of all four genome elements was observed (Fig. 3C). Earlier we have shown that G4 DNA stabilization results in copy number reduction in small plasmid (Mishra et al., 2019). The change in genome copy number has also been reported when the coding sequences of genome segregation proteins were deleted in this bacterium (Maurya et al., 2019a; Maurya et al., 2019b). Change in genome copy number and higher frequency of anucleate cells in Δ*topoIB* mutant suggested an important role of drTopo IB in the segregation genome in *D. radiodurans*.

**Fig 3.**
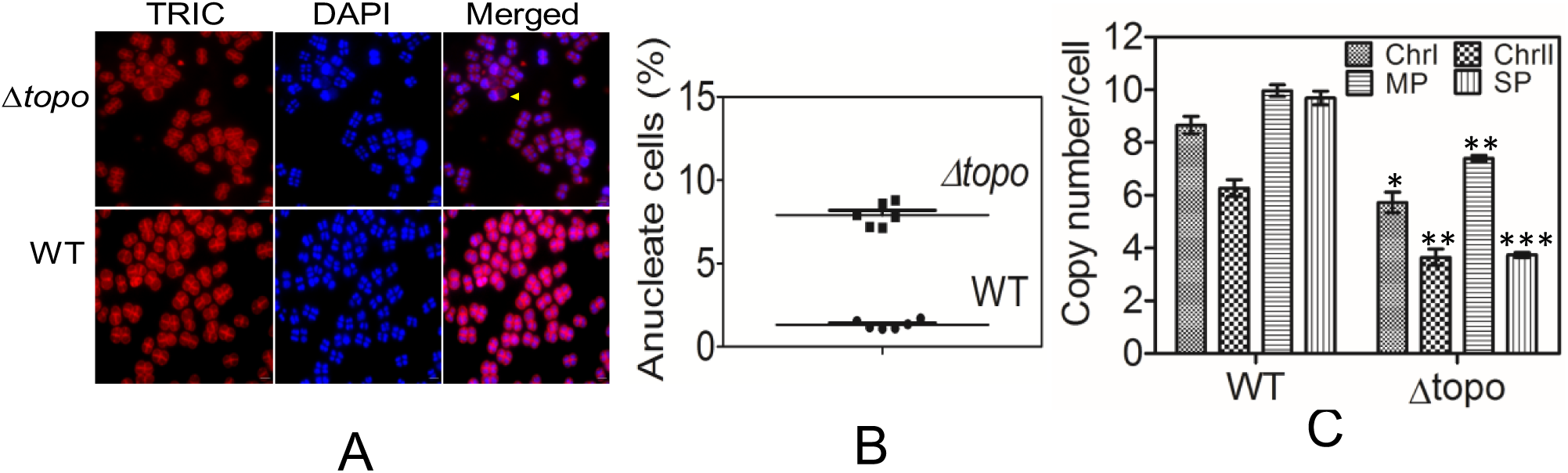
Effect of *topoIB* disruption on nucleoid morphology and stable inheritance of genome in *Deinococcus radiodurans*. The wild type (WT) and *topoIB* (Δtopo) cells were DAPI (DAPI) and Nile Red (TRIC) stained and observed for nucleoid and membrane visualization respectively under microscope (A). The percentage of anucleated cells were calculated through the DAPI staining of the nucleoids in wild type (WT) and *topoIB* cells (Δtopo) (B). Similarly, the *D. radiodurans* wild type (WT) and *topoIB* (Δtopo) cells were grown to log phase and genome copy number of chromosome I (Chr I), chromosome II (Chr II), megaplasmid (MP) and small plasmid (SP) per cell was calculated by quantitative PCR as described in methods (C). The data given are means ± standard errors of the mean (SEM). The Students’s t test was done to see the statistical significance of the genome copy numbers of different elements. Ns (not significant), P > 0.05; **, P<0.01; ***, P<0.001; ****, P<0.0001. The microscopic data shown is the representative picture of three independent experiments done separately.

### 3.3 drTopo IB interacts with genome maintenance proteins in *Deinococcus radiodurans*

Since, drTopo IB disruption showed a significant change in stable inheritance of genome elements in dividing population, we got curious to check the interaction of drTopo IB with proteins involved in segregation and cell division in this bacterium. Physical interaction of genome with proteins in genome segregation was studied by immunoprecipitation and Bacterial Two Hybrid (BACTH) system. For BACTH analysis, the coding sequence of topo IB, genome partition proteins like ParB1, ParB2, ParB3 and DNA replication initiation protein (DnaA) were cloned in BTH vectors pUT18 and pKNT25 for their expression of these proteins as fusion with T18 and T25 domains of CyaA, respectively. *E. coli* BTH101 co-expressing topo IB with above genome partition proteins in various combinations were monitored for CyaA regulated β-galactosidase expression. Results showed that topo IB interacts with all the tested genome partition proteins ParB1, ParB2 and ParB3 as observed by the reconstitution of CyaA mediated β-galactosidase expression (Fig 4A). For co-immunoprecipitation *E. coli* strain BL21 (DE3) pLysS cells expressing His-topoIB through pETtopoIB were co-transformed with ParB1, ParB2, ParB3 and DnaA proteins with T18 tag. The transformants were checked for interaction. The results from co-IP analysis of cell free extract made from the cells co-expressing His-topoIB and T18 tagged genome partition proteins also confirmed their interactions *ex vivo* (Fig 4B). Besides genome segregation proteins, drTopo IB also interacted with DnaA. However drTopo IB interaction was not observed with cell division protein FtsZ.

**Fig 4.**
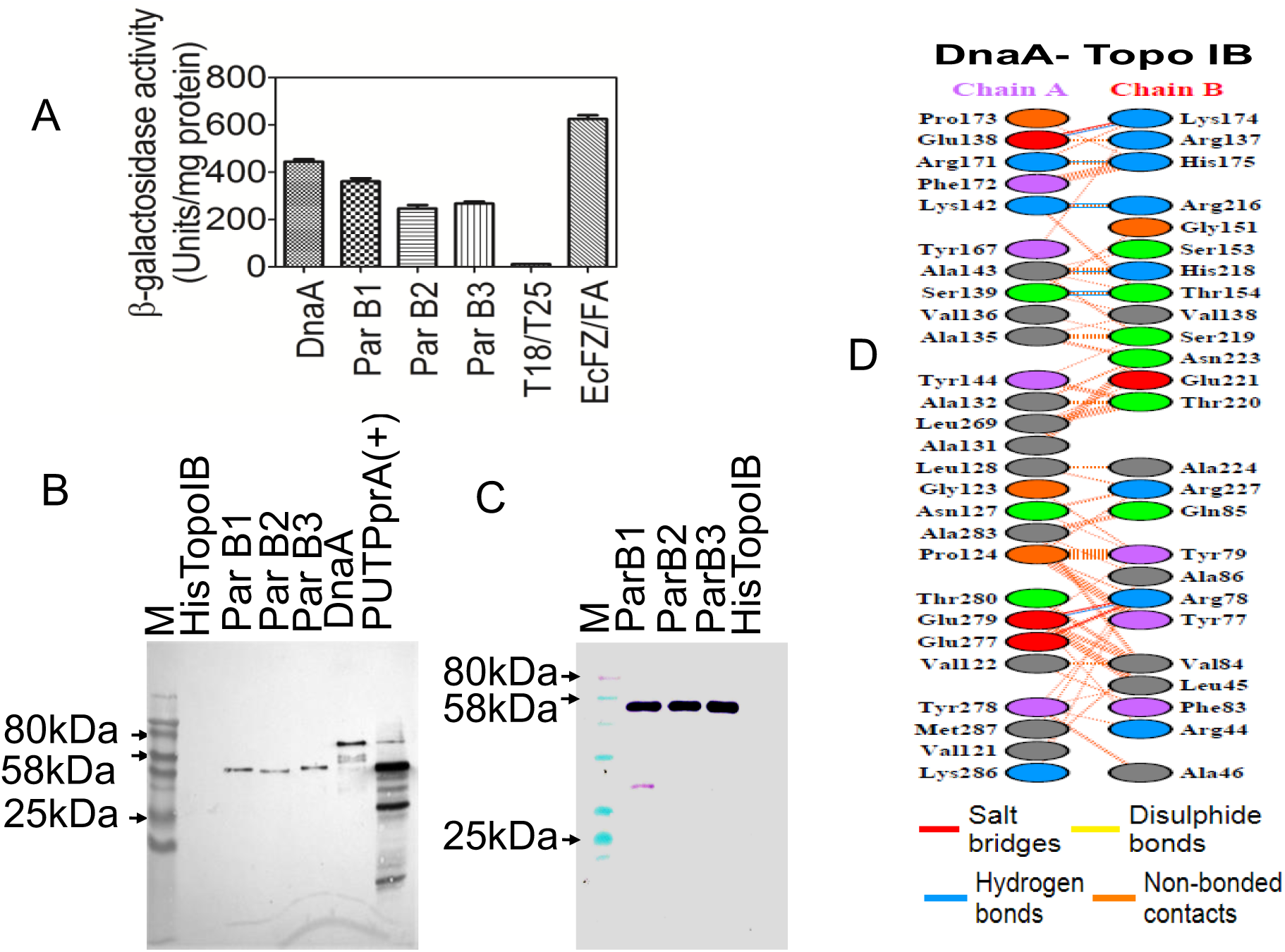
drTopo IB interacting partners in *Deinococcus radiodurans*. The wild type cells co-expressing drTopo IB on pUT18/PKNT25 (pUTtopoIB, pKNTtopoIB) with one of each ParB1, ParB2, ParB3 and DnaA on pKNT25 were assayed for reconstitution of CyaA regulated β-galactosidase enzyme activity expression (A). *E. coli* strain BL21 expressing His tagged topoisomerase IB and T18 tagged ParB1, ParB2, ParB3 and DnaA were monitored for protein-protein interactions through immunoprecipitation using antibodies against polyhistidine tag. The perspective interacting partner of drTopo IB (topoIB) was detected by immunoblotting with antibodies against T18 domain of CyaA. Cell expressing His-topoIB on pET28+ vector was used as negative control and cells expressing T18 tagged PprA as positive control (B). Interaction of drTopo IB with different ParB proteins in *D. radiodurans* was showed by co-expressing polyhistidine tagged drTopo IB and T18 tagged ParB proteins. The cell free extract was immunoprecipitated with antibodies against polyhistidine tag and blotted with T18 antibodies to detect the interactive partners. The D. *radiodurans* cells expressing only His tagged drTopo IB (HistopoIB) were used as negative control (C). *In silico* analysis of protein-protein interaction was carried out as established interface surfaces between drTopo IB (Topo IB) and DnaA as described in methods. The putative amino acids that are interacting with each other are shown (D).

The interaction of drTopo IB with these proteins was also checked in *D. radiodurans*. For that the His-topoIB was co-expressed on low copy number plasmid pRadhistopo with T18 tagged ParB1, ParB2 and ParB3 genome segregation proteins in the bacterium *D. radiodurans*. The cell-free extract of the co-transformants were immunoprecipitated with antibodies against polyhistidine tag and precipitated proteins were checked for the presence of different genome segregation proteins using T18 antibodies. As shown, the ParB proteins were co-immunoprecipitated with drTopo IB (Fig 3C). Further *in-silico* protein–protein interaction studies were performed for drTopo IB and DnaA. A significant number of hydrogen bonds (8 bonds) between drTopo IB and DnaA were predicted, which supported our results (Fig 3D). The above results suggest that drTopo IB physically interacts with ParB and DnaA proteins in this bacterium.

### 3.4 drTopo IB disruption affects cellular dynamics of PprA during post irradiation recovery in *Deinococcus radiodurans*

PprA plays an important role in DNA repair, genome segregation and cell division in *Deinococcus radiodurans* (Narumi et al., 2004; Kota et al., 2016; Devigne et al., 2016). Earlier, it has been shown that PprA regulates the activities of both drTopo IB and DNA gyrase and its localization patterns on the nucleiod mark the growth phase in cells (Kota et al., 2014b; Devigne et al., 2013). Therefore, we got curious to study the localization pattern of PprA in *topoIB* mutant grown under normal as well as gamma irradiated conditions. We observed that GFP-PprA localizes either on mid cell nucleiod (MCN) or on the septum trapped nucleoid (STN) depending upon growth phase in wild type cells grown under normal conditions (Fig 5A & 5B). However, after 2h post irradiation recovery period (PIR), most of the wild type cells had GFP-PprA foci on septum trapped nucleiod (Fig 5C). These observations agreed with our earlier findings on PprA localization dynamics during PIR (Kota et al., 2014b). However, in *topoIB* mutant, the majority of the cells showed foci in the periphery of the nucleiod (Fig 5D & 5E). Besides these, *topoIB* cells also showed different pattern of GFP-PprA localization during PIR when compared with wild type. For instance, at 2h PIR most of the wild type cells had GFP-PprA foci on STN while majority of *topoIB* cells had PprA on MCN (Fig 5F). This might suggest that the absence of drTopo IB can alter the cellular dynamics of PprA during normal and gamma radiation stressed conditions in *D. radiodurans*. This observation was compared with 2 more proteins ParA3 and DivIVA having different physiological functions (like ParA3 is involved in genome segregation while DivIVA is involved in cell pole determination in rod-shaped bacteria) and the results were insightful. For instance, the localization of ParA3 was affected but not DivIVA in the absence of drTopo IB in *topoIB* cells (Fig. S3). Both PprA and ParA3 proteins are involved in genome segregation and absence of drTopo IB in cells had resulted in altered cellular localization pattern of these proteins. These findings together reveal that drTopo IB plays a key role in stable inheritance of genome elements in dividing cells, most likely by affecting the cellular dynamics of genome segregation proteins including PprA in *D. radiodurans*.

**Fig 5.**
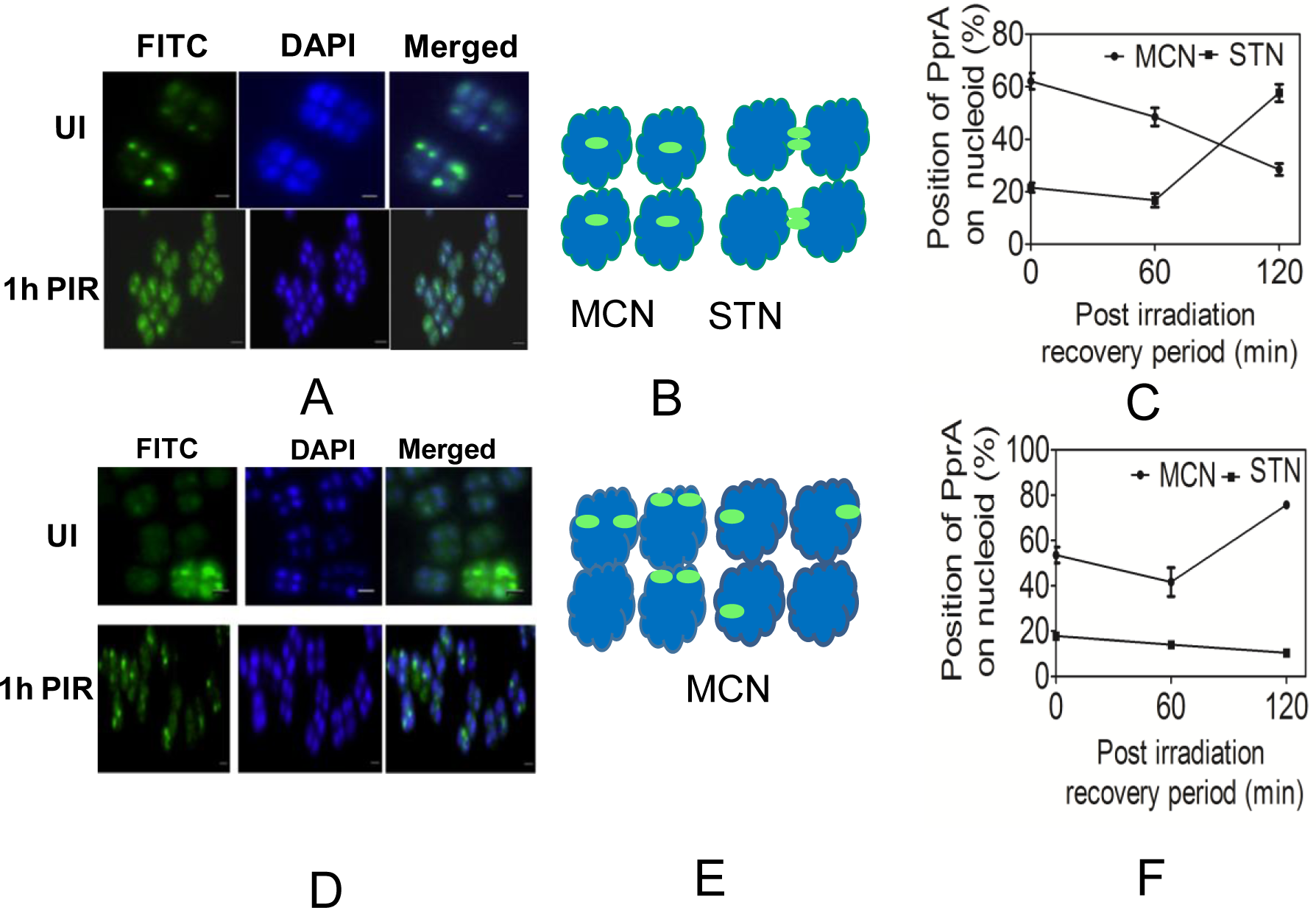
Effect of c on cellular localization of PprA in *Deinococcus radiodurans*. The wild type *D. radiodurans* (A, B, C) and *topoIB* (D, E, F) cells expressing GFP-PprA were grown under normal (UI) and gamma stressed (1h PIR) conditions. These cells were stained with DAPI and imaged under DAPI and FITC channels (A, D). The cellular localization of the GFP-PprA protein on mid cell nucleiod (MCN) and septum trapped nucleiod (STN) was schematically represented in wild type (B) and Δ*topoIB* (E). The percentage GFP-PprA foci on MCN and STN at 0, 60 and 120 mn PIR was represented in wild type (C) and in *topo IB* (F) cells. The Microscopic data shown are the representative pictures of three independent experiments done separately.

### 3.5 drTopo IB localization on nucleoid is cell division stage dependent

The *D. radiodurans* cells expressing drTopo IB-RFP under *groESL* constitutive promoter on low copy plasmid were examined microscopically. Results showed that the protein forms foci on the nucleoid (Fig 6A). Careful examination of a large number of cells showed that the position of drTopo IB localization on the nucleoid is not fixed and more than one foci per nucleoid was observed in large number of cells. Further analysis of these images revealed the dynamic association of this protein with the mechanism underlying nucleoid segregation prior to the septum formation (Fig S4). We also noticed the existence of arbitrary drTopoIB-RFP protein on septum/ring especially in the cells where the genome duplication was completed (Fig 6A). This got further clarified by staining the septum of cells expressing drTopoIB-RFP with vancomycin. We observed drTopoIB localization is perpendicular to the new septum formation and is parallel to the duplicated genome movement (Fig 6B). When we expressed ParA3-RFP and DivIVA-RFP from *groESL* constitutive promoter in *D. radiodurans*, we didn’t observe drTopoIB localization pattern for these proteins. This suggested that drTopoIB localization on the division septum is characteristic to drTopoIB and might suggest its role in genome segregation. Topoisomerase enzymes play the very important role during resolution of intertwined duplicated circular chromosome and this function is performed by topoisomerase IV and tyrosine recombinases in *E. coli* (Usongo et al., 2013). *D. radiodurans* genome lacks topoisomerase IV while DNA gyrase does both supercoiling and decatenation functions on DNA (White et al., 1999; Kota et al., 2016; Devigne et al., 2016). Topoisomerase IB enzyme is related to tyrosine recombinases family proteins and relax both positive and negative supercoiled DNA (<u>Krogh</u> and <u>Shuman,</u> 2002). The drTopoIB localization on nucleiod, interaction with genome segregation proteins and foci formation in perpendicular to new septum and parallel to old septum conceptually indicate drTopoIB role in genome segregation. Further, the results also suggest that drTopoIB localization on nucleoid depends upon the cell division state of the cells as it localizes perpendicular to new septum in actively growing cells and parallel with old septum in resting cells or cells preparing for cell division.

**Fig 6.**
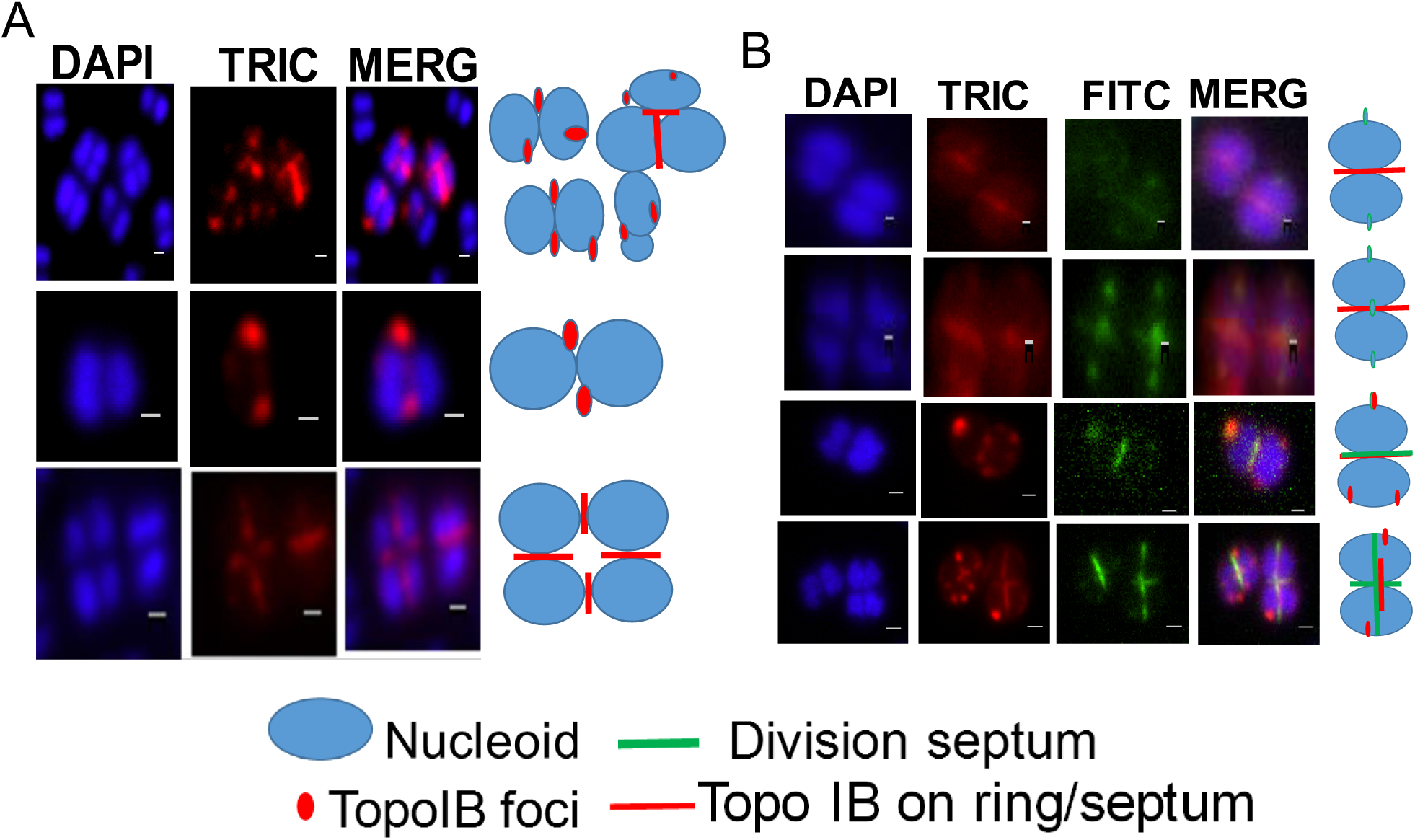
Cellular localization of drTopo IB with respect to septum in *Deinococcus radiodurans*. The wild type *D. radiodurans* cells expressing drTopoIB-RFP (topoIB-RFP) from a constitutive *groESL* promoter were stained with DAPI and vancomycin-fluorescent-BIODPY (Van-FL) and imaged under TRIC, DAPI, FITC channels for protein, nucleoid and membrane imaging respectively. Data shown here is the representative of mixed population (mixed), diplococcic (Diplococci) and tetrad (Tetrad) cells depicting the probable localization of drTopoIB-RFP in cells stained with DAPI (A) and DAPI and vancomycin-fluorescent-BIODPY (B) DAPI stained images were merged (MERG) with FITC and TRIC images as case may be, and their schematic representations with respect to nucleoid and septum are shown beside each panel.

## 4. Discussion

*Deinococcus radiodurans* is a Gram-positive bacterium with multipartite genome system having two chromosomes and 2 plasmids and each of these elements are present in 4-10 copies per cell (White et al., 1999). The GC rich genome of this bacterium is densely packed with guanine repeats (G motifs) having signature for G4 DNA structure formation. Many such G motifs have been shown to form G4 DNA structure (Beaume et al., 2013; Kota et al., 2015a). Recombinant purified drTopo IB has been shown to destabilize G4 DNA structure *in vitro* (Kota et al., 2015b). Here we have brought forth some evidence to suggest the *in vivo* function of drTopoIB in the regulation of gene expression and G4 DNA structure dynamics in *D. radiodurans*. This function of drTopoIB could be through its topoisomerase activity where this enzyme could make topology dependent nick in G4 DNA structure or by changing supercoiling characteristics of dsDNA or both.

Topoisomerases role in DNA metabolism associated to genome integrity have been well documented in both bacteria and eukaryotes. Earlier, the regulatory role of topoisomerases in the non-canonical DNA structure formation and gene expression has been suggested possibly by relaxing the superhelicity of dsDNA. For instance Top I, a type IB enzyme prevents the accumulation of R-loops, shows high affinity binding to G4 DNA and promotes the formation of intermolecular G4 structures (<u>Kim</u> and Robertson 2017, Arimondo et al., 2000). In *Saccharomyces cerevisiae*, the topoisomerases role in gene expression regulation has been shown by maintaining the chromatin state for transcription initiation (Pedersen et al., 2012). Similarly, the Top I role has also been reported in the transcription of genes required during embryogenesis in *Caenorhabditis elegans* (Lee et al., 2001). In *Mycobacterium tuberculosis*, both topoisomerase I and DNA gyrase were shown localizing with RNAP, to take care of transcription induced torsional stress (Ahmed et al., 2017). These findings together might suggest the drTopo IB role in regulation of gene expression by altering the topological structure of promoter or non-canonical secondary structure of the DNA in the regulatory regions.

Interestingly, we also observed an otherwise redundant role of drTopo IB in genome maintenance where the stable inheritance of genetic materials was affected in *topoIB* cells. These cells showed increased number of anucleated cells and a significant population of cells showed the diffused nucleoids. Since, bacterial chromosome segregation is an active process and requires coordination between many proteins involved in DNA replication, segregation of duplicated genome and cell division processes, this role of drTopo IB was not surprising but shown for the first time in this bacterium. In *E. coli*, TopoIII and TopoIV are involved in decatenation of duplicated genomes while topoisomerase I acts indirectly by influencing the supercoiling state of the genome (Seol et al., 2013; Zechiedrich et al., 2000). In *Mycobacterium smegmatis* Topo I is detected at the *ter* region of the genome and implicated its role in genome segregation through decatenation of duplicated genomes (Rani and Nagaraja, 2019). In *D. radiodurans*, the genome partitioning proteins were characterized and their roles in chromosome segregation have been shown. Earlier, the role of PprA protein in genome segregation and cell division was demonstrated through its interaction with DNA topoisomerases. Here, we show that drTopo IB interacted with genome partition proteins and absence of drTopo IB could affect PprA dynamics on the nucleoid during PIR as well as ParA3 localization in general. Since, the localization pattern of cell division protein FtsZ (Fig 7) and DivIVA (Fig S3) was not effected in *topoIB* cells, drTopo IB role mainly in genome maintenance possibly by unlocking the interlinks between the duplicated genomes and by functional interaction with genome partition proteins could be suggested.

**Fig 7:**
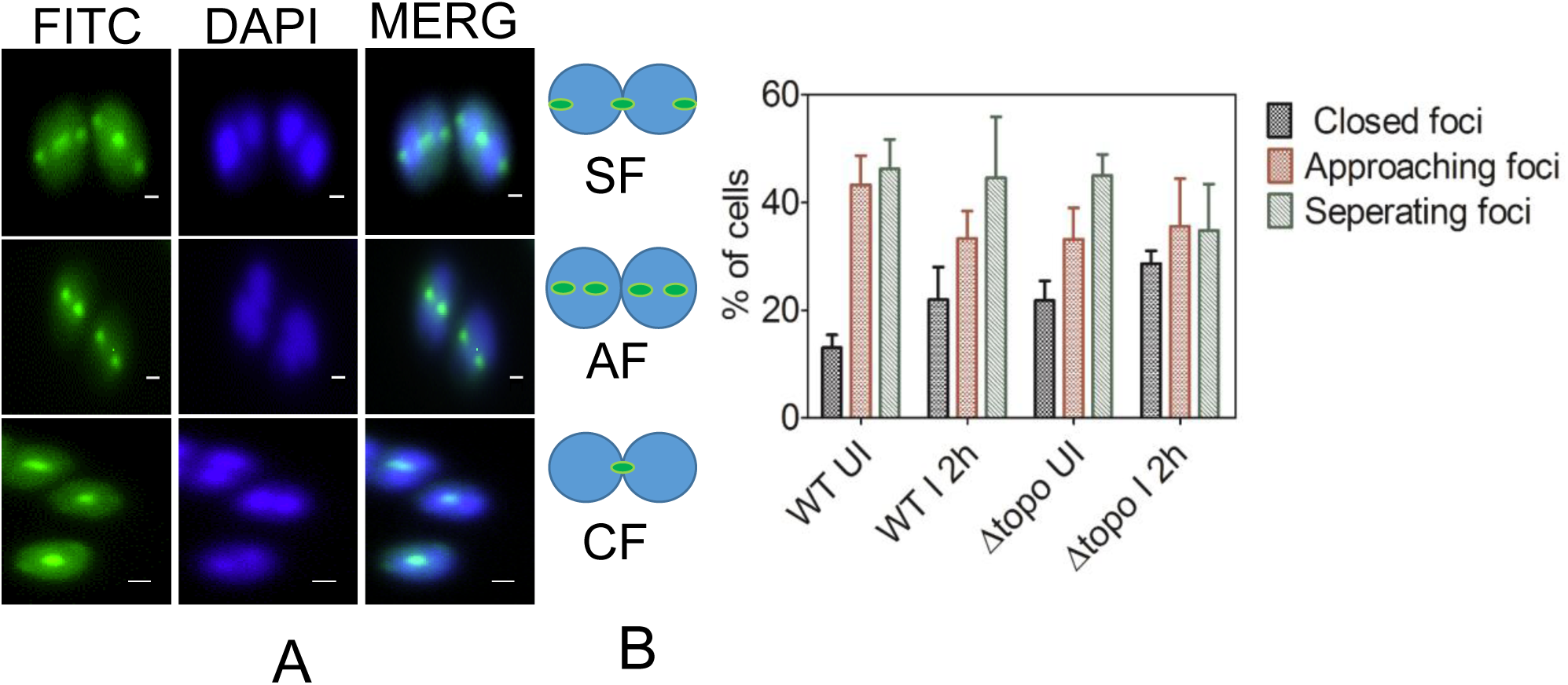
Cellular localization of FtsZ in wild type and *topo IB* cells of *D. radiodurans* grown under normal and gamma stressed conditions. FtsZ protein polymerization pattern in wild type and *topo IB* cells was checked by expressing GFP-FtsZ from pVHSMFtsZ plasmid (A). A heterogeneous population of wild type and *topo IB* cells expressing GFP-FtsZ showing FtsZ foci at juxtaposed position on membrane (separating foci), foci progressed toward each other (approaching foci) and foci merged in the centre (closed foci) as shown schematically (B). Percentage of cells with separating foci (SF), approaching foci (AF) and closed foci (CF) were calculated from unirradiated (UI) and 2h after post irradiated (2HPIR) cells and plotted (C).

The nucleoid in this bacterium majorly forms ring-like or toroidal structures and to some extent crescent and branched rod structures depending on the phase of cell cycle. Recently, it was shown that multiple *oriC* of chromosome I was localized at the periphery while *ter* loci were accumulated mostly at the centre in the nucleoid (Floc’h et al., 2019). Our results on drTopo IB localization revealed a number of drTopo IB-RFP foci on the nucleiod and they are arranged at the peripheral region (Fig. 6A & S4). The drTopo IB interacts with DNA replication initiation protein DnaA, as well as a centromere binding protein ParB and the centromere-like sequences are scattered at multiple locations in chromosome I of *D. radiodurans* (Charaka et al., 2012). Therefore, a possibility of drTopo IB activity around origin of replication and/or *ter* region at least in chromosome I cannot be ruled out and will require independent investigations.

In conclusion, the disruption of *topoIB* from the genome in *D. radiodurans* has resulted in more than one phenotypic change *albeit* all seems to be associated with its known function in DNA metabolism. Furthermore, drTopo IB ability to regulate G4 DNA structure dynamics and promoter activity along with its role in genome maintenance primed a hypothesis that drTopo IB is an important protein in the maintenance of genome structure and function in this bacterium too. Further studies would be required to understand how the activities of other topoisomerases are influenced by each other in unlocking the events required during segregation of multipartite doughnut shaped nucleiod genome, as well as on how these macromolecular complexes are spatially organized at the *oriC* and centromeric-region in this organism. The available results clearly suggest drTopo IB role in regulation of non-canonical DNA structure dynamics and genome maintenance possibly by bringing these two seemingly distant events together through macromolecular interactions.

## Acknowledgements

We acknowledge Ganesh K. Maurya for studies on *in-silico* protein-protein interactions and qPCR experiments. Miss Reema Chaudhary is thankful to the Department of Atomic Energy, Government of India, for research fellowship.

